# Assessment of Genetic Resources to Major Tomato Diseases: Insights from wild germplasm and public *S. lycopersicum* collections

**DOI:** 10.1101/2025.03.04.641566

**Authors:** Ehtisham Hussain, Chien-yu Cheng, I-Min Huang, Chen-yu Lin, Samrin Gul, Ijaz Rasool Noorka, Assaf Eybishitz, Chutchamas Kanchana-udomkan, Maarten van Zonneveld, Ya-ping Lin

**Author notes:** **Corresponding author:** Ya-Ping Lin.

## Abstract

Tomato (*Solanum lycopersicum* L.) production is severely impacted by biotic stresses, leading to significant yield losses. Developing genetically resistant cultivars presents a sustainable alternative to chemical control, which is often costly and ineffective against evolving pathogens. This study utilized molecular markers to assess genetic resistance to key tomato diseases, including Fusarium wilt, late blight, bacterial wilt, root-knot nematode, *Tomato Mosaic Virus* (ToMV), and *Tomato Yellow Leaf Curl Virus* (TYLCV), across 964 tomato accessions comprising both wild germplasm and cultivated varieties. Wild germplasm exhibited the highest frequencies, with *S. peruvianum* identified as a rich reservoir of resistance loci. Markers associated with TYLCV (*Ty-1/3, Ty2*) and Fusarium wilt (I2(OH)) were particularly prevalent in wild species. The potential for gene pyramiding potential was evident, with some wild accessions carrying up to six resistance loci. In contrast, resistance loci were limited in *Solanum lycopersicum*. Two *S. lycopersicum* germplasm collections were screened: a geographically representative collection from four Southeast Asian national genebanks and a genetically representative core collection from global public genebanks. The highest resistance frequencies in *S. lycopersicum* were observed for bacterial wilt-associated markers SLM12-2 and SLM12-10. However, the overall scarcity of resistance alleles in *S. lycopersicum* emphasizes the need for further introgression of resistance genes from wild relatives. This study provides valuable genetic insights into tomato germplasm for combating biotic stress, forming a foundation for sustainable breeding strategies to enhance disease resistance and safeguard global tomato production.

## Introduction

Tomato (*Solanum lycopersicum* L.) is one of the most widely cultivated vegetable crops globally, valued for both fresh consumption and processed food products. In 2022, global tomato production reached approximately 186.8 million metric tons, spanning 4.9 million hectares (FAOSTAT, 2024). However, tomato production is continuously challenged by biotic stresses, with over 200 pests and pathogens affecting yield and fruit quality (Bai et al., 2018). These include fungi, bacteria, viruses, and nematodes, which not only reduce productivity but also impact the crop’s marketability and profitability (Panno et al., 2021). The threat is particularly severe in tropical and subtropical regions, where favorable conditions enable the proliferation of destructive diseases such as tomato yellow leaf curl disease (TYLCD), bacterial wilt (*Ralstonia solanacearum*), and late blight (*Phytophthora infestans*) (Hanssen et al., 2010; Fry, 2008; Navas-Castillo et al., 2011). Additional threats include *tomato spotted wilt virus* (*TSWV*), *tomato mosaic virus* (*TMV*), root-knot nematodes (*Meloidogyne* spp.), and Fusarium wilt (*Fusarium oxysporum* f. sp. *lycopersici*) (Mahfouze et al., 2022; Chitwood-Brown et al., 2021; Lizardo et al., 2022). The widespread impact of these diseases underscores the urgency of implementing sustainable strategies to enhance disease resistance in tomato cultivars.

The genetic diversity of cultivated tomatoes has been significantly reduced due to domestication bottlenecks and selective breeding, which have prioritized yield and fruit quality over resistance traits (Tamburino et al., 2020). To overcome this limitation, wild *Solanum* species serve as invaluable reservoirs of resistance genes. Species such as *S. pimpinellifolium* and *S. pennellii* have contributed resistance to multiple pathogens (Bohra et al., 2022). Although introgressing resistance traits from wild relatives into cultivated tomatoes requires extensive backcrossing, past breeding efforts have demonstrated its feasibility and effectiveness (Migicovsky & Myles, 2017). Notably, breeding programs in the Netherlands have expanded the genetic diversity of modern tomato cultivars by eightfold since the 1960s, largely due to the strategic incorporation of wild-derived resistance traits (Schouten et al., 2019). This increased genetic base has significantly improved both resilience and phenotypic variability.

In addition to wild relatives, cultivated tomato varieties, including commercial hybrids and landraces, offer valuable genetic resources for breeding programs. While commercial hybrids are developed primarily by private breeding initiatives and require trait stabilization for long-term use, landraces maintained in public genebanks exhibit genetic diversity and resilience to various environmental stresses. Landraces have shown particular promise in providing resistance to late blight and other pathogens, sometimes outperforming commercial cultivars, particularly in organic farming systems (Maxim et al., 2022; Boziné-Pullai et al., 2021). However, fully utilizing these resources remains challenging due to the complexity of disease resistance phenotyping and the vast number of tomato accessions in public genebanks worldwide.

Public genebanks, such as the World Vegetable Center (WorldVeg), maintain extensive collections of tomato landraces and crop wild relatives (Bauchet & Causse, 2012; Ebert & Chou, 2014). Despite considerable progress in characterizing these genetic resources, two critical challenges persist. First, breeding programs often utilize a limited subset of available germplasm, highlighting the need for broader collaboration and resource-sharing to enhance genetic diversity. Second, comprehensive resistance data remain incomplete, necessitating the development of cost-effective and scalable screening strategies to efficiently identify promising accessions.

For over half a century, the WorldVeg has been at the forefront of tomato breeding for tropical regions, recently integrating marker-assisted selection (MAS) into its pre-breeding efforts to address prevalent diseases. This study builds upon that foundation by systematically screening and documenting disease resistance-related genetic diversity within a representative tomato germplasm collection. By incorporating MAS and DNA fingerprinting into genebank databases, this initiative aims to enhance breeding efficiency, reduce reliance on labor-intensive phenotyping, and provide a valuable resource for sustainable disease resistance breeding. Furthermore, the generated data will serve as a decision-support system to optimize genebank management and regeneration strategies, ultimately contributing to the development of more resilient tomato cultivars.

## Materials and Methods

### Plant material

A total of 964 tomato accessions were evaluated for the presence of genetic resistance alleles against various diseases (Supplementary Table 1). These accessions were categorized into three distinct groups: the wild tomato collection, the Southeast Asian *S. lycopersicum* collection, and the global *S. lycopersicum* core collection. The first group contained 371 wild tomato accessions representing nine *Solanum* species; they were maintained at the WorldVeg Genebank. The second group consisted of 291

**Table 1.**
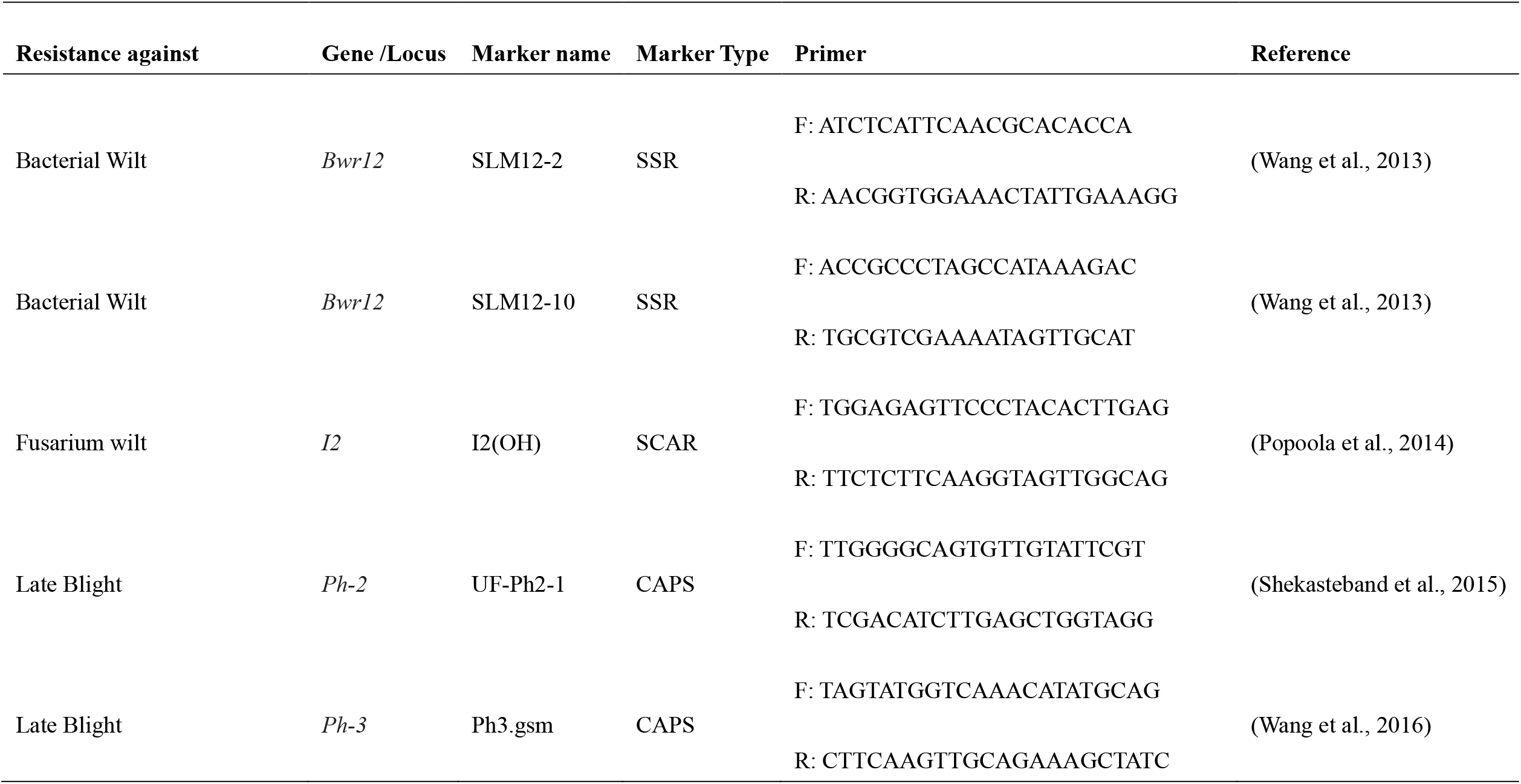
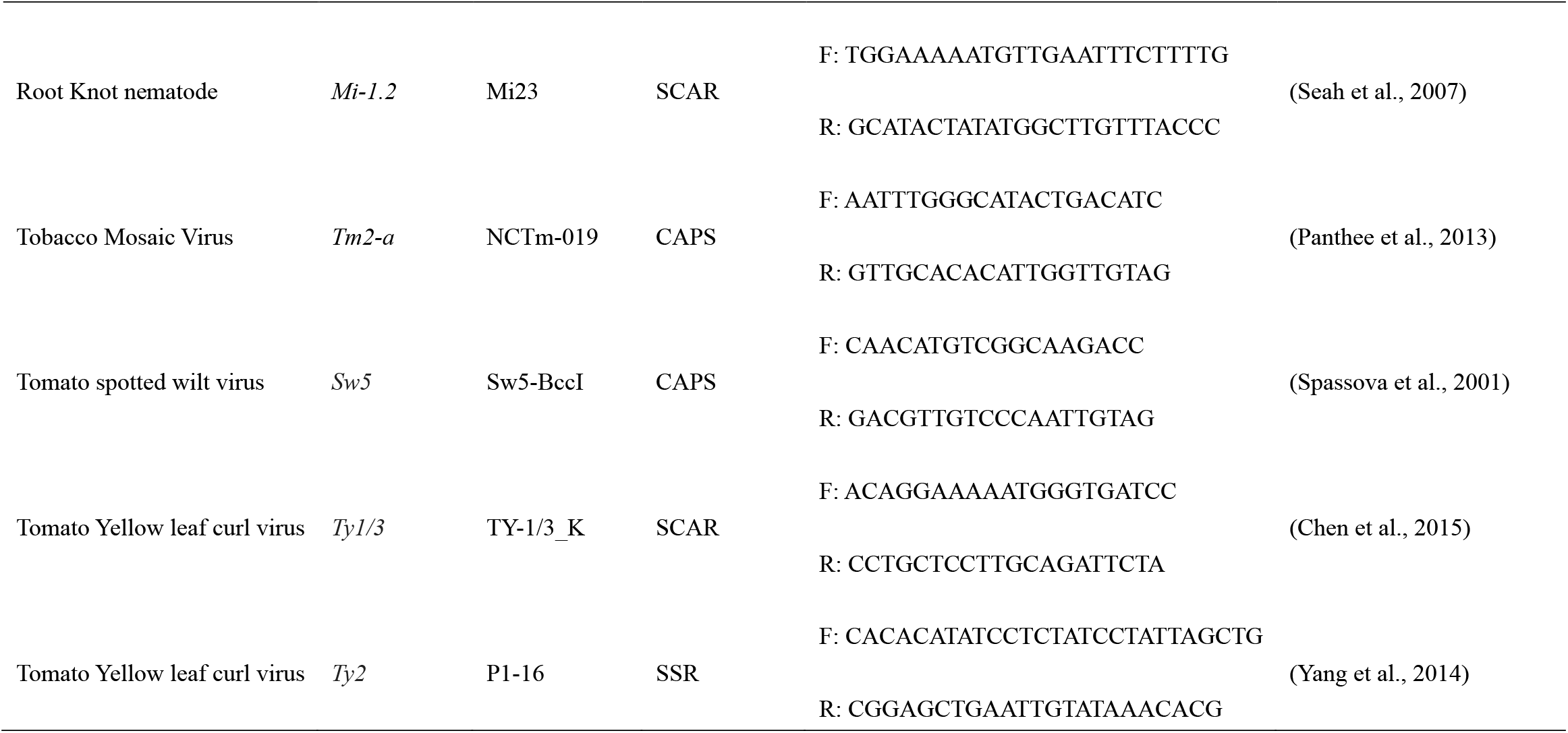
Disease-resistant molecular markers used in the study.

*S. lycopersicum* accessions collected from four Southeast Asian countries, Malaysia, Thailand, the Philippines, and Vietnam, preserved at the WorldVeg Genebank and recently repatriated to their countries of origin. This subset was designed to represent *S. lycopersicum* diversity in Southeast Asia. The third group included 302 accessions from a genetically representative core collection of cultivated tomatoes, excluding wild accessions (Barchi et al., 2019).

### DNA preparation

All plant materials, along with 11 resistant and susceptible check lines, were cultivated in a greenhouse environment. Young leaf tissues were harvested for genomic DNA extraction following the cetyltrimethylammonium bromide (CTAB) protocol (Tanksley, 1995). The quality and concentration of the extracted DNA were assessed using a Qubit® 2.0 Fluorometer (Invitrogen). DNA samples were diluted to a final concentration of 20 ng/μL and stored at -20°C for future use.

### PCR preparation

Ten molecular markers were employed for DNA fingerprinting to identify disease-resistant quantitative trait loci (QTLs) (Table 1). Polymerase chain reaction (PCR) was performed in 96-well plates, with a total reaction volume of 15 μL per well. The reaction mixture consisted of 10.1 μL of nuclease-free water, 1.5 μL of 10X buffer containing 15 mM MgCl_2_, 0.8 μL of dNTP mix (Cyrusbioscience, Inc.), 0.5 μL of forward and reverse primers (10 μM concentration), 0.1 μL of Taq polymerase (Thermo Fisher Scientific), and 2 μL of DNA template.

### PCR conditions

PCR amplifications were carried out using a Bio-Rad S1000™ thermal cycler (Bio-Rad, USA). The cycling conditions for all sequence-characterized amplified region (SCAR) and simple sequence repeat (SSR) markers were as follows: an initial denaturation at 95°C for 10 minutes, followed by 35 cycles of denaturing at 95°C for 30 seconds, annealing at 55°C for 45 seconds, and amplifying at 72°C for 45 seconds. A final extension was performed at 72°C for 5 minutes, with an additional cooling step at 15°C for 5 minutes.

For cleaved amplified polymorphic sequence (CAPS) markers, the PCR cycling conditions as follows: an initial denaturation at 95°C for 10 minutes, followed by 35 cycles of denaturing at 95°C for 30 seconds, annealing at 55°C for 1 minute, and amplifying at 72°C for 2 minutes. A final extension was conducted at 72°C for 5 minutes, followed by a cooling step at 15°C for 5 minutes. Following PCR, restriction enzymes (*Hae*III, *Hin*fI, *Hin*cII, and *Bcc*I) (New England BioLabs® Inc.) were used to digest PCR products corresponding to the markers NCTm-019, UF-Ph2-1, Ph3-Gsm1, and Sw5 markers, respectively. The digestion reactions were prepared in a 10 μL volume containing 3.9 μL of nuclease-free water, 1 μL of 10X CutSmart™ Buffer (New England BioLabs® Inc.), 0.1 μL of the restriction enzyme, and 5 μL of the PCR product. Digestion reactions were incubated overnight at 37°C to ensure complete enzyme activity.

### Gel electrophoresis

PCR and digestion products were separated using a 6% polyacrylamide gel in 0.5X TBE buffer. Electrophoresis was conducted at 135V for 35–40 minutes, depending on fragment size. Gels were stained with SYBR Safe dye and visualized using a Bio-1000F gel imager (Microtek, Taiwan). A 50 bp DNA ladder (Bio-50bp™ DNA Ladder, Protech Technology Enterprise Co., Ltd.) was included as a molecular size reference for fragment length determination.

## Results

### Distribution of resistance alleles across the *Solanum* species

The application of 10 PCR-based molecular markers successfully identified resistance, susceptible and heterozygous alleles, with a missing rate of only 1.49% across the analyzed tomato germplasm (Fig. 1). A comprehensive assessment of 371 wild tomato germplasm accessions revealed substantial variation in the distribution of resistance alleles associated with major tomato diseases (Fig. 2). The frequency of resistance alleles varied across markers, with I2(OH), linked to the *I2* for Fusarium wilt resistance, exhibiting the highest frequency (55.8%). This was followed by markers associated with *Tomato Yellow Leaf Curl Virus*; P1-16, associated with the *Ty2* QTL (34.0%), and TY-1/3_K, associated with *Ty1/3* (31.8%). In contrast, the bacterial wilt marker SLM12-10, linked to the *Bwr12* QTL, displayed the lowest frequency (4.3%), followed by the late blight markers Ph3.gsm (6.2%) and UF-Ph2-1 (6.5%).

**Figure 1.**
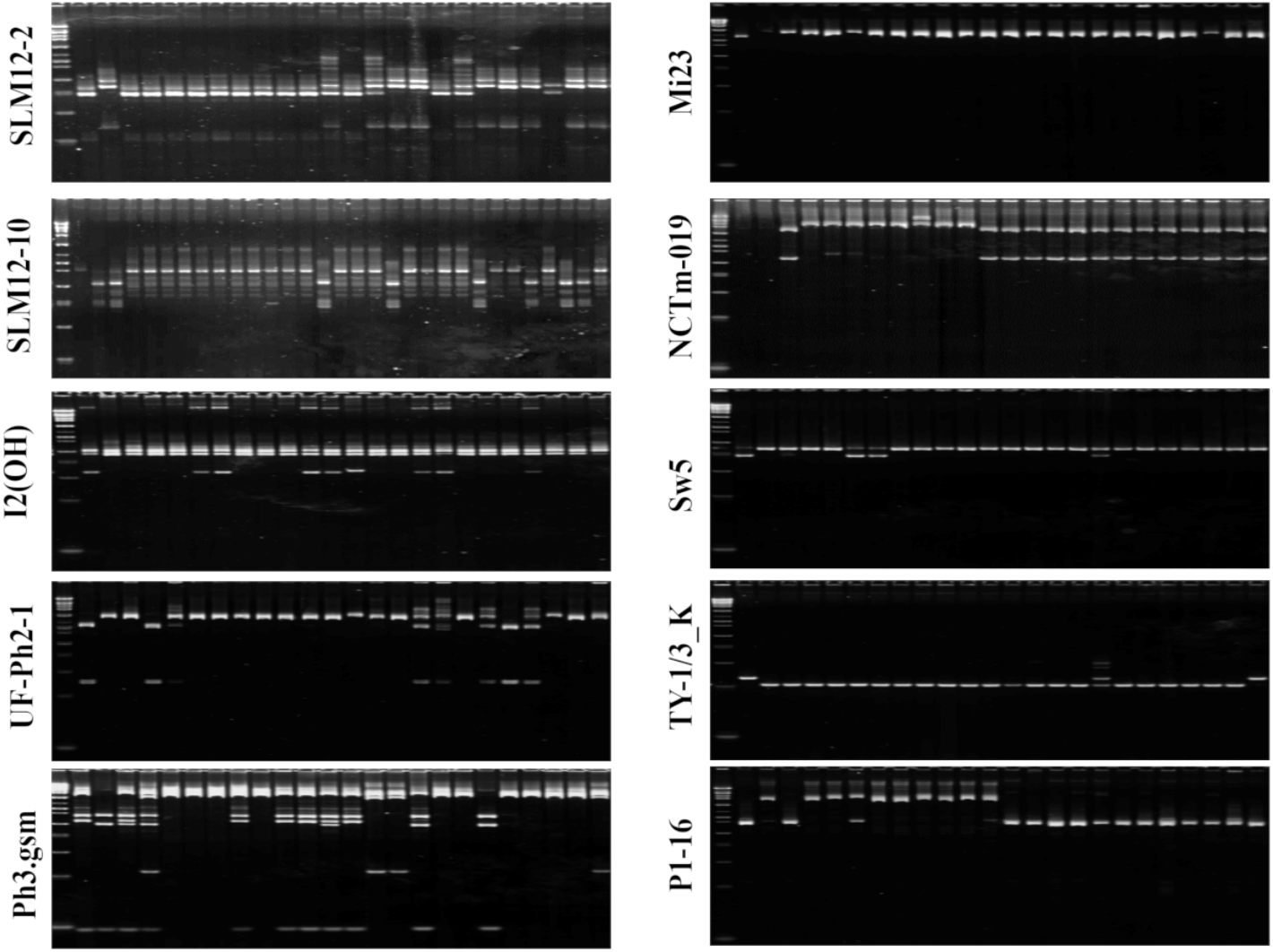
Results of polyacrylamide gel electrophoresis for the molecular markers in this study. From left to right of all the visualized gels consisted of: L1 50bp DNA ladder, L2 resistant check, L3 susceptible check. For NCTm-019, the resistant and susceptible checks were absent in this gel; for Mi23, susceptible check was absent in this gel.

**Figure 2.**
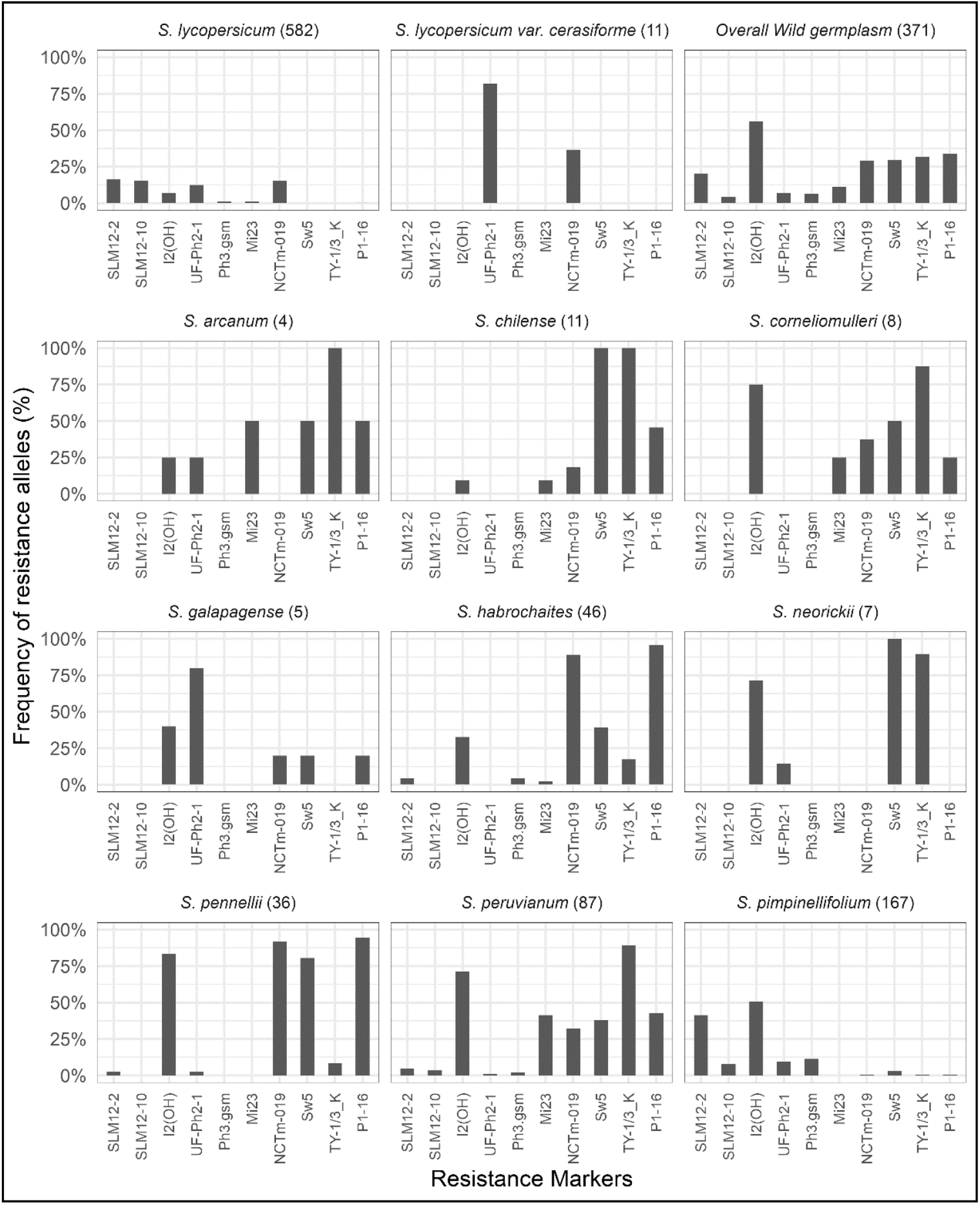
Frequency of ten resistance markers among the *Solanum* species in this study.

The wild tomato germplasm analyzed encompassed nine distint *Solanum* species, each displaying unique resistance allele distributions. Notably, *S. chilense* exhibited a 100% frequency for resistance alleles associated with *Ty-1/3* and *Sw5*, while the Ty2 allele was present at a frequency of 45.5%.

Additionally, the *Tm2-a, I2*, and *Mi-1.2* resistance QTLs were detected at frequencies of 18.2% and 9.1%, respectively. Among the 167 accessions of *S. pimpinellifolium*, resistance alleles were predominantly detected for I2(OH) (50.9%) and SLM12-2 (41.3%), while other markers were observed at lower frequencies, including Sw5 (3%), Mi23 (0%), UF-Ph2-1 (9.6%), Ph3.gsm (11.4%), TY-1/3_K (0.6%), P1-16 (0.6%), NCTm-019 (0.6%), and SLM12-10 (7.8%).

*S. peruvianum* exhibited the highest frequencies for TY-1/3_K (89.7%) and I2(OH) (71.3%), alongside notable frequencies for P1-16 (41.7%), Sw5 (38.1%), and NCTm-019 (32.1%). Similar trends were observed in *S. habrochaites* and *S. pennellii* for eight of the ten resistance markers; however, *S. pennellii* exhibited higher frequencies of Sw5 and I2(OH) compared to *S. habrochaites*. Interestingly, four species, *S. chilense, S. cornelimulleri, S. galapagense*, and *S. neorickii*, lacked resistance alleles for markers Ph3.gsm, SLM12-2, and SLM12-10 in all analyzed accessions (Fig. 3). These findings underscore the species-specific diversity of resistance alleles within wild tomato germplasm and highlight their potential as valuable genetic resources for breeding disease-resistant cultivars.

**Figure 3.**
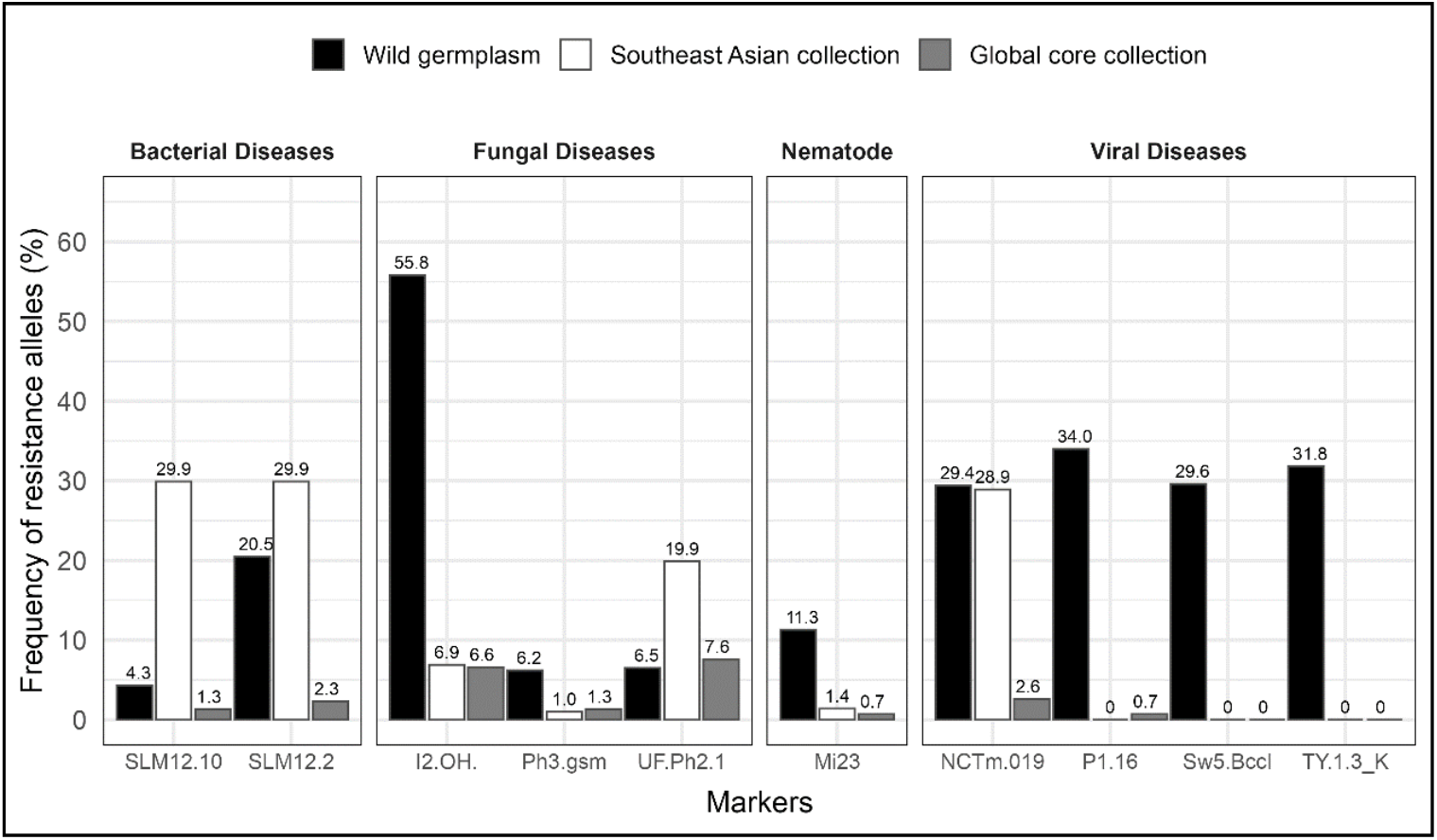
Comparisons of the frequency of resistance alleles among the wild germplasm, the Southeast Asian collection, and the global core collection regarding to various diseases.

### Genetic resistance in the geographically representative *S. lycopersicum* collection in Southeast Asian

This gene pool consisted of 291 *S. lycopersicum* accessions collected from four Southeast Asian countries: Malaysia, Thailand, the Philippines, and Vietnam. Across the entire collection, the highest allele frequencies were observed for the bacterial wilt markers SLM12-2 and SLM12-10 at 29.9%, followed by the *Tm2-a*-associated marker NCTm-019 at 28.87%. Between the late blight resistance loci, UF-Ph2-1 exhibited a higher frequency (19.93%) than Ph3.gsm (1.03%). Notably, markers Sw5, TY-1/3_K, and P1-16 were completely absent in this Southeast Asian *S. lycopersicum* collection. The Mi23 marker, conferring root-knot nematode resistance, was rarely detected, with frequencies of 0% in Vietnam and the Philippines, 2.73% in Thailand, and 3.23% in Malaysia.

A country-specific analysis revealed significant variation in allele distribution. Among the 31 Malaysian accessions, the bacterial wilt resistance markers SLM12-10 and SLM12-2 exhibited the highest frequencies, at 51.61% and 45.16%, respectively. Resistance loci associated with Sw5, Ph3.gsm, TY-1/3_K, and P1-16 were absent. In Thailand, among 110 accessions, the highest frequency was observed for NCTm-019 (72.73%), followed by UF-Ph2-1 (27.27%) and I2(OH) (13.64%). The Philippines, representing the largest subgroup with 146 accessions, exhibited the highest frequencies for SLM12-10 and SLM12-2 (43.15% each). while markers associated with all four viral resistance loci (Sw5, TY-1/3_K, P1-16, and NCTm-019) were entirely absent. In Vietnam, four accessions displayed a 100% frequency for NCTm-019, with SLM12-2 and SLM12-10 each present at 50%.

### Genetic resistance in the genetically representative *S. lycopersicum* collection

The genetic assessment of 302 *S. lycopersicum* accessions from the global tomato core collection revealed limited genetic resistance to disease. Only 62 accessions (20.53%) carried at least one resistance allele. Among the evaluated markers, UF-Ph2-1 exhibited the highest resistance frequency at 7.62%, followed by I2(OH) at 6.62%. Markers Mi23 and P1-16 were detected at identical frequencies (0.66%), while Sw5 and TY-1/3_K were entirely absent across the collection. Notably, resistance associated with the *Ph2*, which confers late blight resistance, was more prevalent than the *Ph3*. Overall, genetic resistance within this collection was minimal.

### Resistance frequency across the genetic resources

A comparative analysis among the wild germplasm and the two *S. lycopersicum* collections revealed significant variation in resistance allele frequencies. The Sw5 and TY-1/3_K markers, absent in *S. lycopersicum*, were present in the wild germplasm at frequencies of 29.6% and 31.8%, respectively (Fig. 3). Similarly, I2(OH) and P1-16 were more frequent in the wild germplasm. Notably, four markers, Bwr12-linked markers, Ph2, and NCTm-019, demonstrated substantially higher resistance frequencies in the Southeast Asian collection compared to the global core collection. Conversely, six markers, I2(OH), Ph3.gsm, Mi23, Sw5, TY-1/3_K, and P1-16, exhibited only marginal differences between these two collections. The global core collection displayed the lowest overall resistance, with UF-Ph2-1 showing the highest frequency at just 7.62% (Fig. 3). These findings underscore the importance of wild germplasm and the Southeast Asian collection as reservoirs of genetic resistance for breeding programs.

### Concurrence of resistance alleles

The distribution of accessions carrying resistance alleles varied significantly across wild germplasm and *S. lycopersicum*. A total of 50 wild accessions (13.5%), 72 *S. lycopersicum* accessions (24.7%) from Southeast Asia, and 240 *S. lycopersicum* accessions (79.5%) from the core collection lacked resistance alleles. Conversely, accessions carrying only one of the nine evaluated resistance alleles were observed at frequencies of 18.3% in the wild germplasm, 38.5% in the Southeast Asian collection, and 17.9% in the core collection (Figure 4). The wild germplasm displayed the highest diversity in the concurrent presence of multiple resistance alleles (Table S1). Specially, three wild accessions contained six resistance alleles, 18 contained five, and 91 exhibited three (Table 2). Among *S. peruvianum* accessions, 13 out of 87 (15%) carried more than five resistance alleles, highlighting its potential as a valuable resource for resistance breeding.

**Figure 4.**
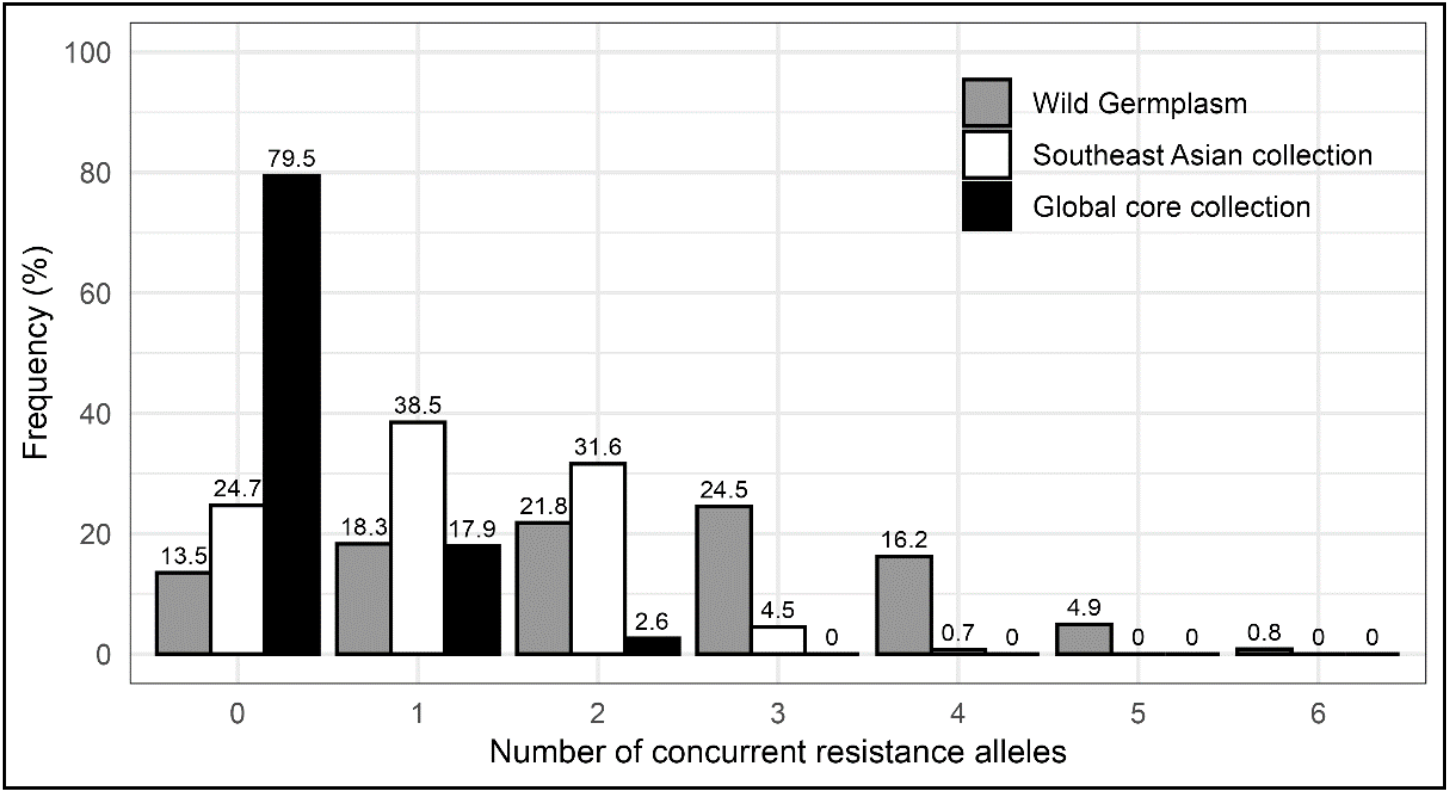
Frequency of concurrent resistance alleles among wild germplasm, the Southeast Asian collection, and the global core collection.

**Table 2.**
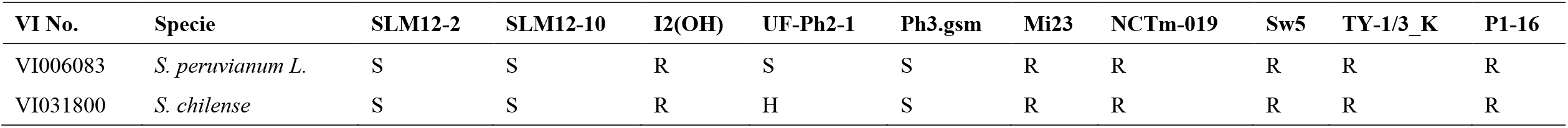
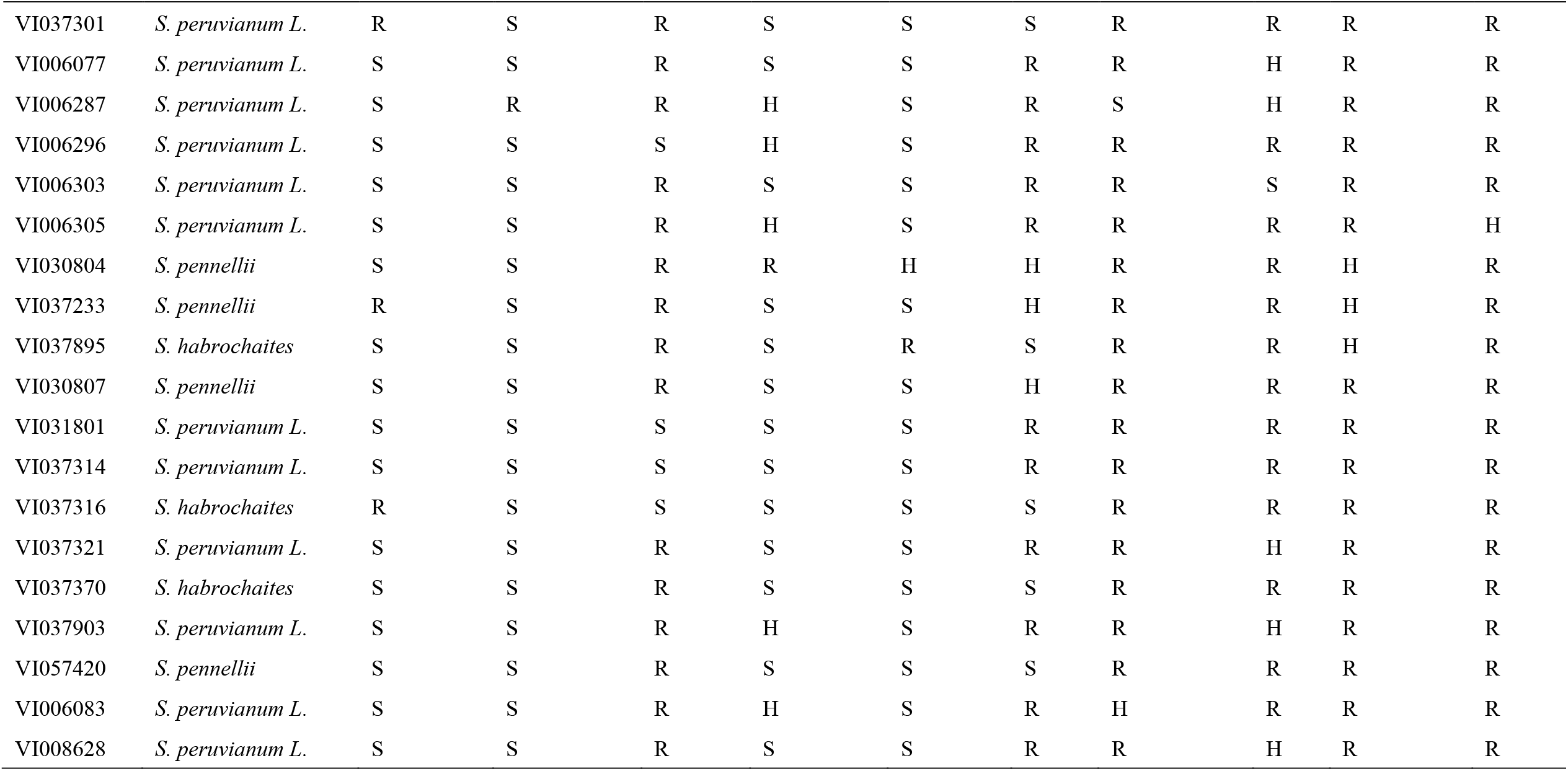
Resistance-allele DNA fingerprints of the 21promising accessions.

In contrast, the Southeast Asian collection exhibited a lower incidence of multiple resistance alleles, with two accessions carrying four resistance alleles. However, accessions with two or more resistance alleles frequently included those carrying both markers for the *Bwr12* locus, which accounted for 66.4% of multiple-resistance accessions in this collection. These results emphasize the genetic richness of the wild germplasm and the Southeast Asian collection in harboring diverse resistance traits, underscoring their potential for breeding disease-resistant cultivars, particularly in tropical regions.

### Discussion

Tomato production faces significant yield losses due to biotic stresses, particularly diseases. Chemical control measures, though widely used, are often costly, environmentally hazardous, and ineffective against certain pathogens. Overuse of these chemicals raises concerns about food safety and sustainability. In contrast, cultivars with inherent genetic resistance provide a more sustainable alternative, reducing chemical dependency while ensuring stable yields under disease pressure. Molecular markers have become essential in breeding programs for disease resistance, enabling precise selection of single resistance genes and gene pyramids (Lee et al., 2015).

In this study, molecular markers were employed to evaluate resistance to key tomato pathogens, including those responsible for Fusarium wilt, root-knot nematode, late blight, bacterial wilt, *TMV*, and *TYLCV*. The results revealed significant variation in genetic resistance among the wild germplasm, the Southeast Asian collection, and the global core collection, with the latter displaying the most limited resistance. This limitation may stem from historical breeding practices that prioritized yield and agronomic traits, inadvertently selecting against resistance alleles (Derbyshire et al., 2024). Similar trends have been reported in previous studies, such as Tiwari et al. (2023), who documented the absence of *Mi-1* for root-knot nematode resistance in tomato varieties, and Şeker et al. (2023), who found *Ph-2* absent in diverse tomato lines. The successful introgression of resistance traits from the wild relatives, such as *Mi-1* from *S. peruvianum* (Messeguer et al., 1991) and *Ph-1, Ph-2*, and *Ph-3* from *S. pimpinellifolium* (Merk et al., 2012), has historically played a crucial role in improving disease resistance in cultivated tomatoes. These findings reaffirm the significance of wild germplasm as a vital source of resistance alleles for breeding programs.

Beyond the wild relatives, the Southeast Asian collection exhibited substantially greater genetic diversity compared to the global core collection, likely due to its heterogeneous nature (Byrne et al., 2020). This diversity results from centuries of natural and unconscious human selection, during which landraces have co-evolved with biotic and abiotic stresses, accumulating beneficial alleles for adaptability and disease resistance (Sánchez-Martín et al., 2017; Casañas et al., 2017). The broader genetic base of this collection enables adaptation to diverse environments and traditional farming systems (Dwivedi et al., 2016; Fullana-Pericàs et al., 2019). Consequently, the Southeast Asian accessions represent an invaluable genetic resource for breeding resilient tomato varieties (Maxim et al., 2022).

MAS has proven instrumental in reducing labor and costs associated with phenotype-based disease screening. Co-dominant markers, such as SCAR markers, have demonstrated efficacy in tracking resistance to bacterial speck, Fusarium wilt, *TMV*, and *TSWV*, facilitating precise allele selection in breeding programs (Zhang & Panthee, 2021). This study underscores the potential of MAS in pyramiding multiple resistance alleles, thereby enhancing durability against evolving pathogens. For example, combining *Ty-2, Ty-3, Ph-2*, and *Ph-3* has successfully yielded tomato lines resistant to both late blight and *TYLCV* (Kaushal et al., 2024). The synergistic effect of *Ph-2* and *Ph-3* in conferring enhanced late blight resistance has been well-documented, with significantly higher resistance observed in genotypes harboring both alleles (Arafa et al., 2017; Kumar et al., 2022). Resistance gene pyramiding thus offers a strategic approach to mitigating the declining effectiveness of individual resistance genes while countering emerging virulent pathogen strains. (Foolad et al., 2008; Fry, 2008). While molecular markers provide a more efficient alternative to traditional phenotypic assessments, their reliability depends on the strength of their linkage to target resistance traits (Collard & Mackill, 2008; Hasan et al., 2021). Discrepancies between marker presence and actual resistance phenotypes can result from environmental influences and epistatic interactions (Kumar et al., 2024; Sharma et al., 2022; Zabotina, 2013; Arens et al., 2010). This limitation highlights an important future direction: the need for field validation of resistance phenotypes. While molecular markers provide valuable insights into genetic resistance, their effectiveness must be confirmed through field trials under natural disease pressure. Future studies should integrate molecular marker analysis with phenotypic evaluations across multiple locations and seasons to ensure the practical utility of identified resistance traits.

A key challenge in genebank management is the vast genetic diversity within collections, often comprising thousands of accessions with unknown or poorly characterized traits. This study addresses that challenge by generating resistance marker data to facilitate more precise trait-based searches in genebanks. Molecular annotation of accessions enhances resource accessibility and expedites the development of disease-resistant crop varieties (Ebert & Schafleitner, 2015). Additionally, molecular markers can help identify promising accessions for breeding programs, ensuring valuable resistance alleles are not lost during seed multiplication. Moreover, integrating resistance marker databases with genebank management systems promotes global collaboration and data sharing. Publicly accessible platforms enable researchers to contribute to germplasm characterization, creating a continuous feedback loop that enhances genetic resource conservation and utilization. Such initiatives ensure that rare and novel resistance alleles are preserved and effectively incorporated into breeding efforts, further supporting the development of resilient tomato cultivars suited to diverse agroecological conditions.

### Conclusion

This study assessed genetic resistance alleles for major diseases affecting tomatoes in tropical regions, analyzing both wild germplasm and cultivated varieties. The results highlight that wild germplasm harbors significantly higher resistance levels than cultivated tomatoes, making it an invaluable resource for breeding programs. While some cultivars carry resistance alleles, their low frequency emphasizes the need for ongoing efforts to identify and integrate resistance sources into breeding pipelines. The findings also underscore the importance of molecular data for genebank management, facilitating the efficient conservation and utilization of genetic resources in breeding applications. Although transferring resistance traits from wild relatives to commercial cultivars presents challenges, advancements in molecular breeding offer promising strategies to overcome these barriers. These technologies enhance precision in resistance gene introgression, paving the way for the development of resilient tomato varieties that support sustainable agriculture in disease-prone environments.

## Acknowledgements

This work was supported by grants from the National Science and Technology Center, Taiwan (MOST 110-2313-B-125-001-MY3), awarded to Y.-P. Lin and the Ministry of Foreign Affairs of Taiwan (Taiwan Asia Vegetable Initiative). We also thank the long-term strategic donors to the World Vegetable Center, including the governments of Taiwan, Germany, Thailand, the Philippines, South Korea, Japan, UK, USAID, and ACIAR.

## Author contribution

Ya-ping Lin conceptualized and supervised the research. Ya-ping Lin and Maarten van Zonneveld secured the funding for the project. Ehtisham Hussain, Chien-yu Chen, I-Min Huang, and Chen-Yu Lin performed the experiments. Ehtisham Hussain performed the data curation, statistical analyses and prepared the figures. The first draft of the manuscript was written by Ehtisham Hussain and Ya-ping Lin. All authors commented on previous versions of the manuscript. All authors read and approved the final manuscript.

## Supplementary data

Supplementary Table 1 Accession information and the disease-resistant DNA fingerprint in this study.

